# Nanopore sequencing methods detect cell-free DNA associated with MRD and CNS infiltration in pediatric Acute Lymphoblastic Leukemia

**DOI:** 10.1101/2021.09.27.462067

**Authors:** Shilpa Sampathi, Yelena Chernyavskaya, Meghan G. Haney, L. Henry Moore, Isabel A. Snyder, Anna H. Cox, Brittany L. Fuller, Tamara J. Taylor, Tom C. Badgett, Jessica S. Blackburn

## Abstract

Acute Lymphoblastic Leukemia (ALL) patients that are positive for minimal residual disease (MRD) after therapy or have leukemic infiltration into the central nervous system (CNS) are considered high risk and receive intensive chemotherapy regimens. Current methods to diagnose MRD and CNS infiltration rely on detecting leukemic cells in patient samples using pathology, flow cytometry, or next-generation sequencing. However, leukemic blasts may persist in the patient but not be physically present in bone marrow biopsy or biofluid sample, leading to inaccurate or delayed patient diagnosis. We have developed a nanopore sequencing workflow to detect B-ALL-associated cell-free DNA (cfDNA) in blood and cerebrospinal fluid (CSF) samples. Quantitation of B-cell specific VDJ recombination events in cfDNA samples defined B-ALL clonal heterogeneity. This workflow allowed us to track the response of individual B-ALL clones throughout treatment. Detection of cfDNA also predicted the clinical diagnosis of MRD and CNS disease. Importantly, we identified patients diagnosed as CNS negative who had low B-cell derived cell-free DNA levels in their CSF sample that correlated with B-cell clones present in the bone marrow. These data suggest that cfDNA assays may be useful in detecting the presence of ALL in the patient even when blasts are not in the biofluid sample. Nanopore analysis of cell-free DNA is a simple, rapid, and inexpensive assay that can serve as a valuable complement to traditional clinical diagnostic approaches for ALL.

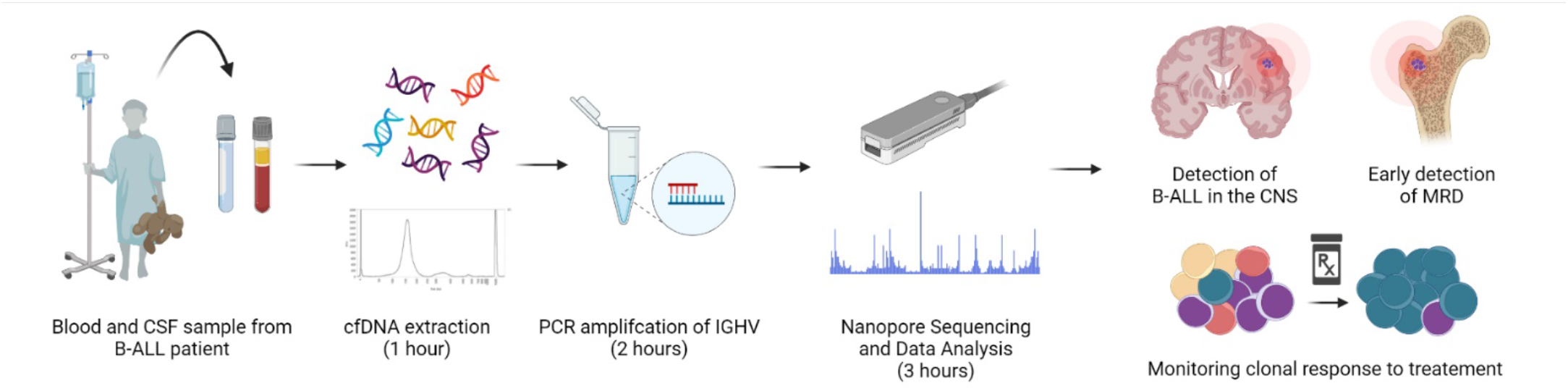

## Introduction

Acute Lymphoblastic Leukemia (ALL) is the most common pediatric malignancy. Despite cure rates approaching 90%, relapsed ALL remains the second leading cause of cancer-related death in the pediatric population. Response to therapy, as measured by the detection of minimal residual disease (MRD) of >0.01% at the end of induction, remains the most powerful prognostic indicator of outcome (1-3). The central nervous system (CNS) is an important site of involvement in ALL, and is a common site for ALL relapse (4, 5). The prevalence of CNS disease in ALL is sufficiently high that every ALL patient receives prophylactic treatment involving multiple doses of intrathecal chemotherapy, which can have adverse and long-term side effects in pediatric patients. Better stratification of patients based on risk status and potential for relapse or CNS disease will rely on more precise diagnostics for ALL.

Current standard clinical methods to diagnose MRD and CNS disease in ALL rely on pathology and flow cytometric detection of blasts in patient samples. Developing more sensitive assays that depend on the molecular detection of leukemic blasts via next-generation sequencing (NGS) has proven difficult for this disease. ALL has one of the lowest mutational burdens of any cancer (6, 7), which precludes the design of targeted, cancer-specific sequencing probes commonly used in NGS assays in other types of cancers. Recently, Adaptive Biotechnologies developed the ClonoSEQ assay, an NGS assay that sequences VDJ rearrangements within T and B cell receptors instead of cancer-specific mutations (8, 9). This strategy is ideal for lymphocytic leukemias, as every T and B cell undergoes VDJ recombination to generate functional B and T cell receptors; the specific sequence of the rearrangement in the receptor is unique to an individual B or T cell. ALL results from the abnormal expansion of typically 1-5 B or T cell clones, each of which will have a unique VDJ sequence (10, 11). ClonoSEQ identifies clones based on a significant expansion of that particular VDJ sequence within the patient’s blood and bone marrow samples at leukemia diagnosis. At the end of treatment, ClonoSEQ can detect as few as 1 in a million cells within a patient sample that carry that same VDJ rearrangement to sensitively and accurately detect MRD. ClonoSEQ is now in a clinical trial in the US for detection of MRD in leukemias and has been implemented in Europe for MRD detection in B-cell lymphoid malignancies (12-14)

The major limitation of the ClonoSEQ method is its reliance on the collection of genomic DNA from leukemia cells within the patient sample. A significant concern for any cell-based assay is the accurate detection of CNS disease in ALL. Research using animal models showed that ALL cells often become embedded in the CNS tissue or attached to spinal nerves (15, 16). Autopsies of ALL patients that succumbed to disease demonstrated that 85% had CNS disease present in the brain or spinal cord at autopsy (17). However, typically only 11% of patients are clinically diagnosed as CNS disease positive based on detection of lymphoblasts in the cerebrospinal fluid (CSF) (4, 5). These findings highlight a critical need to develop more sensitive assays to complement cell-based pathology, flow cytometry, and ClonoSEQ assays, especially pertaining to ALL spread into the CNS.

The utilization of cell-free DNA as a cancer biomarker is transforming how patients are diagnosed and monitored for cancer progression. Cell-free DNA is constitutively released by cancer cells and is found free-floating in patient biofluids (18, 19). Tumor-specific cfDNA increases as tumors grow. The half-life of cfDNA is only 15-120 minutes, making it a valuable and proven biomarker for detecting cancer development in healthy individuals and monitoring tumor response to treatment in cancer patients (20). In 2020, FoundationOne’s Liquid CDx cfDNA test was FDA-approved to detect *EGFR, BRCA1/2, ALK, ATM*, and *PIK3CA* mutations in cfDNA in ovarian, breast, prostate, and lung cancers to stratify patients into treatment regimens (21). Detection of tumor-specific VDJ rearrangements in cfDNA is used to monitor lymphoid lymphoma progression (22-24), and emerging evidence in myeloid malignancies suggests that cfDNA monitoring will be a useful diagnostic tool for leukemia patients as well. For example, in Acute Myelogenous Leukemia, cfDNA assays detected bone marrow relapse 30 days earlier than the standard flow cytometry-based methods (25). Finally, emerging evidence suggests that cfDNA may be more beneficial than cell-based assays in monitoring CNS tumors. For example, in leptomeningeal carcinomatosis, cancer-derived cfDNA was detectable in cases where microscopy did not reveal malignant cells in the CSF, and cfDNA fluctuated over time correlating with CNS tumor burden (26). A comprehensive examination of several types of CNS malignancies showed that cfDNA isolated from the CSF was much more reliable than genomic DNA isolated from cells within the CSF in identifying tumor-associated mutations (27). In this study, cells could not be reliably obtained from CSF samples, and DNA quality from cells was often not suitable for NGS. These data suggest that cfDNA can identify CNS-associated cancers much more accurately than assays that rely on the presence of cancer cells within the CSF. To our knowledge, cfDNA has not been used to examine MRD status or CNS disease in ALL.

Our study outlines a method for isolation and detection of leukemia-derived cfDNA in the peripheral blood and CNS of pediatric B-ALL patients. We developed a simple and inexpensive workflow based on Nanopore MinION sequencing of PCR amplified VDJ rearrangements in cfDNA, which allowed us to analyze as little as 25 picograms of cfDNA per patient biofluid sample. We found that cfDNA can provide a more accurate assessment of B-ALL heterogeneity, as cfDNA sequencing could detect clones not present in the genomic DNA of bone marrow biopsy samples. Plasma cfDNA samples were used to monitor the response of specific clones to treatment and identified B-ALL clones that lead to MRD. Additionally, the cfDNA sequencing workflow could detect B-ALL associated cfDNA in the CSF of patients that had been clinically diagnosed with CNS disease and in some patients diagnosed as CNS-negative. The clearance of CNS-associated clones was confirmed by loss of cfDNA in the patient CSF sample. In total, our data demonstrate the utility of cfDNA in assessing B-ALL heterogeneity and in monitoring B-ALL progression. The low cost and ease of use make the MinION cfDNA workflow ideal for cfDNA analysis in research settings, and the specificity and sensitivity of the assay in detecting MRD and CNS disease suggests that cfDNA may ultimately be a valuable complement to current cell-based clinical assays like ClonoSEQ.

## Results and Discussion

### Characteristics of B-ALL patients

Pediatric B-cell acute lymphoblastic leukemia (B-ALL) patients were admitted to the Kentucky Children’s Hospital at initial diagnosis of B-ALL and enrolled for sample collection. We retrospectively chose samples for use in this study based on CNS disease and MRD status after initial therapy. In total, our cohort consisted of five patients, out of which one patient (AAL-008) was MRD positive at the end of induction. Patient AAL-004 had a few leukemic blasts noted but was clinically MRD negative as blast counts did not reach the threshold for an MRD diagnosis. Patients AAL-010, 009, and 012 were MRD negative at the end of induction, and patients AAL-008 and 010 were positive for CNS disease at the time of diagnosis. **Table 1** summarizes the complete characteristics of these patients. Genomic DNA was isolated from the bone marrow aspirate collected from each B-ALL patient at the time of diagnosis. Cell-free DNA was isolated from plasma and cerebrospinal fluid (CSF) samples collected throughout treatment.

**Table 1.** Patient Characteristics.

| *Subject ID* | *Blast Count at Diagnosis (\*10<sup>9</sup>/L)* | *WBC Count at Diagnosis (\*10<sup>9</sup>/L)* | *Number of Days to Clear Circulating Blasts* | *EOI MRD Status* | *CNS Disease Status* | *Number of Days to Clear CSF* |
| --- | --- | --- | --- | --- | --- | --- |
| *ALL-004* | 0.57 | 3.33 | 9 | Negative* | Negative | N/A |
| *ALL-008* | 1.53 | 4.79 | 9 | Positive | Positive | 8 |
| *ALL-009* | 14.92 | 19.38 | 11 | Negative | Negative | N/A |
| *ALL-010* | 1.58 | 6.32 | 13 | Negative | Positive | 9 |
| *ALL-012* | 23.88 | 31.84 | 11 | Negative | Negative | N/A |

\*Negative MRD at the threshold of 1/10,000 cells but a few leukemic cells noted
WBC, white blood cell; EOI, end of induction; MRD, minimal residual disease; CNS, central nervous system; CSF, cerebrospinal fluid

### Identification of B cell clones via nanopore sequencing of the recombined IGH gene

We developed a novel experimental workflow that combines PCR of the IGH region with the third-generation Oxford Nanopore MinION sequencer (ONT) (28) to detect B-ALL patient-specific IGH clonal rearrangements and track these clones throughout the patient’s treatment. Ultimately, this method can be applied to genomic DNA isolated from blasts in patient samples or cfDNA collected from patient biofluids. In the first step, we used a commercially available PCR kit to amplify the IGH sequence between the conserved framework region 1 (FR1) and the conserved Joining (J) region, which allows for a complete representation of all variable regions within the IGH gene (29). PCR products were subjected to library preparation to add adapters and barcodes that allow for multiplexing, and then samples were run through the benchtop MinION nanopore sequencer. We used a simple pipeline comprised of freely available software was used to analyze the output files: i) MinKNOW, which operates the sequencer and automatically performs simultaneous sequencing, demultiplexing, and barcode trimming using the built-in Guppy toolkit, ii) Minimap2 to map the merged fastq files to create BAM alignment files, and iii) Featurecounts to determine how many reads align with the IGHV region. We have provided step-by-step instructions for analysis using these tools in the Supplemental Methods. From sample collection to data analysis, the entire process takes less than one working day, costs less than $30 per sample, and can be carried out by users with basic computer skills.

### Limit of input DNA detection on nanopore workflow

The recommended amount of input DNA for IGHV PCR is 2 ng (30). As our ultimate goal was to adapt our workflow towards the detection of cfDNA, we determined the lowest level of input DNA that could be used in IGHV PCR and nanopore sequencing. We tested our workflow on healthy tonsil control DNA, ranging from 10-100 pg. The human IGH region is located on the long arm of chromosome 14 (Chr14, **Figure 1A**), and we used the nanopore generated reads that mapped to this locus over other locations as a readout of PCR specificity and sensitivity. Libraries of IGHV PCR reactions from all DNA concentrations tested had reads that mapped to chromosome 14. Predictably, mapping frequency to Chr14 decreased with decreasing input DNA amount; however, we observed only a slight reduction in total mapped reads after we lowered DNA input to 0.025ng (**Figure 1B**). When we distributed mapped reads across all chromosomes, it became evident that non-specific mapping increased inversely with template amount. The signal to noise was indistinguishable at concentrations below 0.025ng, as reads mapping to Chr14 dropped below the levels of mapping to other chromosomes (**Figure 1C**). Nevertheless, even with the lowest input DNA amount, we could detect reads mapping to the identical IGHV clones identified with the highest DNA input amount (**Figure 1D**), suggesting that sensitivity with nanopore sequencing can be maintained even when specificity decreases. This aspect of nanopore sequencing is essential when working with precious patient samples, such as cfDNA, that are too low to quantify by standard means.

**Figure 1.**
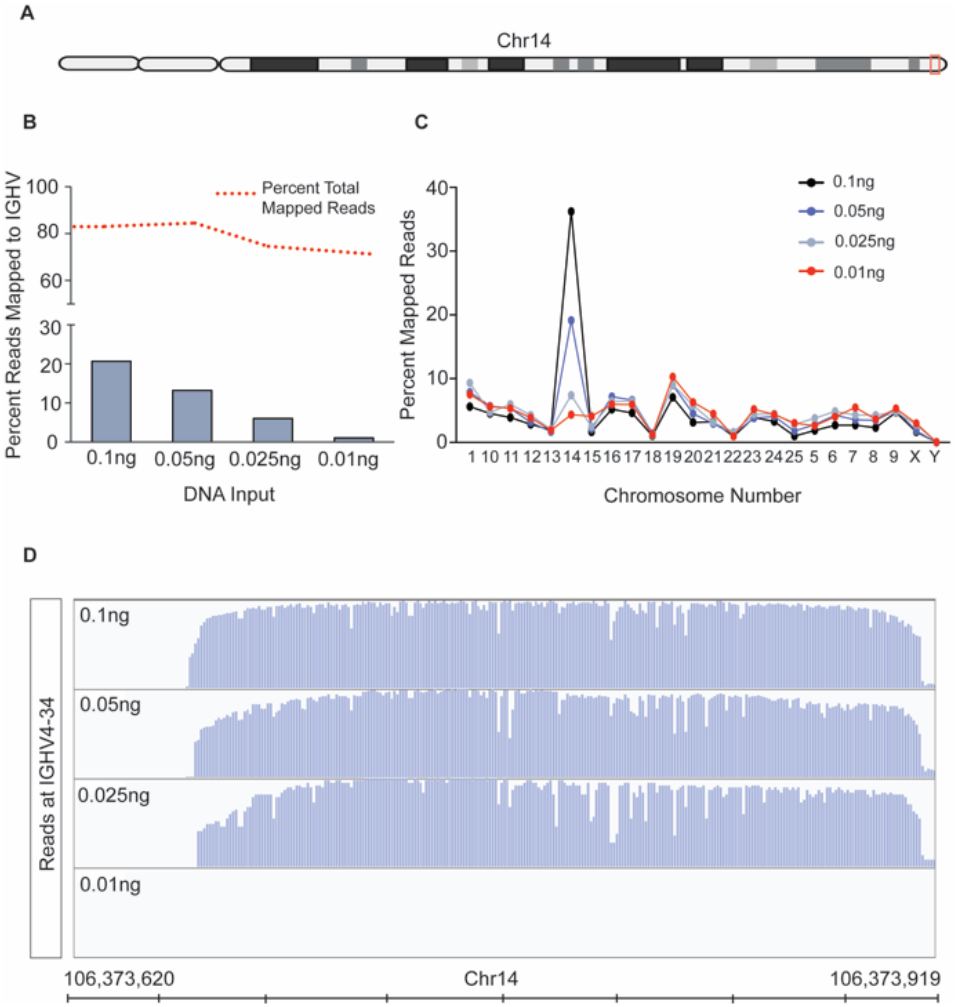
The MinION sequencing workflow can reliably detect IGHV reads in as little as 25 picograms of DNA. A) Schematic of chromosome 14. The red box denotes the region containing the IGHV sequence that all MinION reads were aligned to. **B**) The IGHV region was PCR amplified from the input amount of normal tonsil DNA indicated. Approximately 4,000 reads were obtained from the resulting libraries using nanopore MinION sequencing. The red dashed line shows the percent of all reads that mapped to the human genome, and the bars denote the percentage of those reads that mapped to an IGHV sequence. **C**) The percent of all reads mapped from the samples in (B) are plotted across all chromosomes. **D**) Alignment plots to a selected IGHV, V4-34, shows accurate alignment of reads when input DNA is as little as 0.025ng.

### cfDNA is detectable by the nanopore sequencing workflow

cfDNA is emerging as a robust prognostic biomarker that can be used to assess the tumor heterogeneity and tumor burden (31). Additionally, cfDNA is localized to patient biofluids, so sample collection is less invasive than tumor or tissue biopsy. cfDNA has not previously been assessed in ALL— bioanalyzer traces of cfDNA samples isolated from the plasma of patient AAL-004 and AAL-010 matched expected cfDNA sizes of ∼150 base pairs (**Supplemental Figure 1**). We applied PCR and nanopore sequencing of IGHV to both the genomic DNA (gDNA) isolated from bone marrow aspirate and cfDNA isolated from the plasma at diagnosis. **Supplemental Table 1** details the total number of reads generated from these samples and the percentage mapped to chromosome 14. Only sequences that had an 80% or greater match to the IGHV region of chr14 were considered B cell “clones” and included for further analyses. Clones that comprised ≥ 5% of the total mapped reads were considered significant clones in the sample. We reasoned that a ≥ 5% representation of a particular IGHV sequence in the sample would most likely be due to clonal expansion of the B-ALL. Additionally, we used the same PCR library for nanopore sequencing and MiSeq analysis and detected similar clone identity and clonal abundance across platforms (**Supplemental Figure 2**). These data indicate that the nanopore sequencing workflow can assess the IGH clonal repertoire in B-ALL and identify identical clones in cfDNA. This method is comparable to the more time-consuming and costly next-generation sequencing methodologies routinely used in the clinical setting.

### Nanopore sequencing of cfDNA monitors therapeutic response and MRD

The short half-life of cfDNA makes it an excellent biomarker for assessing leukemia burden and analyzing the response of individual clones to therapy. We analyzed the gDNA from bone marrow biopsy and the cfDNA from plasma collected from 4 patients at diagnosis to identify the expanded clones comprising the B-ALL, as described above. These clones were then tracked in subsequent plasma cfDNA samples to assess the changes in clonal distribution throughout treatment. cfDNA samples were taken ∼ 1 week into induction chemotherapy (denoted as Early-I in **Figure 2**), ∼2 weeks into induction therapy (denoted mid-I), and at the end of induction chemotherapy (denoted EOI). In some cases, we collected cfDNA at the end of consolidation (EOC) therapy, which can occur several months after diagnosis.

**Figure 2.**
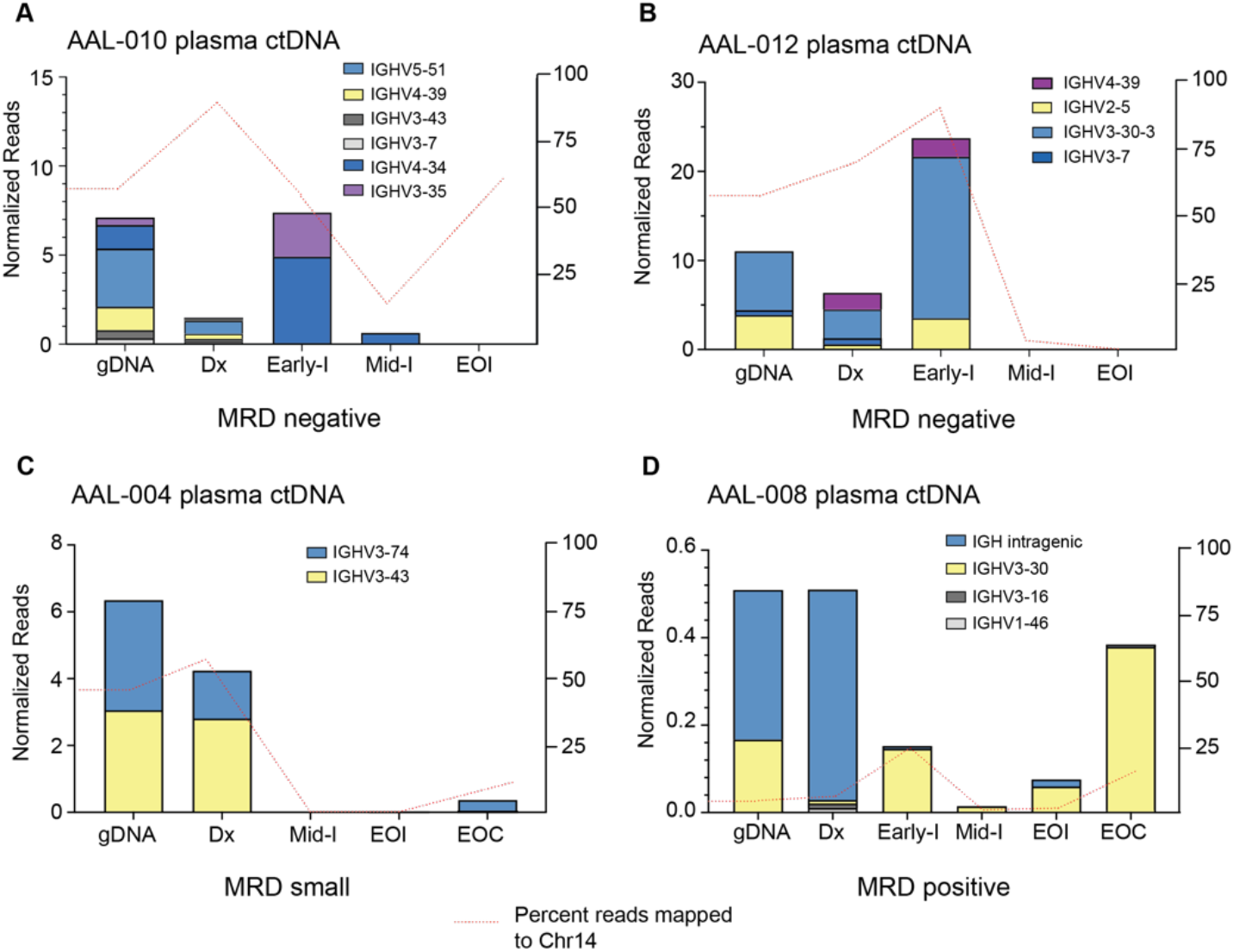
IGHV sequencing of cfDNA can be used to determine B-ALL heterogeneity and to track clonal response throughout the course of treatment. cfDNA was extracted from blood samples of four patients, IGHV regions were PCR amplified, and resultant libraries were sequenced using MinION. Data were analyzed using Galaxy. Clones in the genomic DNA (gDNA) and diagnosis cell-free DNA (Dx) samples that comprised at least 5% of total reads were tracked across all timepoints. Clone abundance is normalized to total mapped reads per library. Percent reads aligned to Chr14 (red line) is a relative indicator of leukocyte abundance at each timepoint/sample. Patients in (**A**) and (**B**) were clinically diagnosed as MRD negative. Patient (**C**) had blasts present in the sample but did not reach the more than 1 blast per 10,000 cell threshold for MRD diagnosis. (**D**) was diagnosed as MRD positive.

Two patients, AAL-010 and AAL-012, were clinically diagnosed as MRD negative at the end of induction therapy. At their initial disease diagnosis, both patients had heterogenous B-ALL, with 6 (AAL-010) or 4 (AAL-012) distinct IGHV sequences present as ≥ 5% of the total mapped reads. In patient AAL-010, we observed a clonal population, IGHV3-35, present in the cfDNA at diagnosis and early induction that was not in the gDNA biopsy sample. All clones were eliminated after treatment, consistent with the clinical diagnosis of MRD-free (**Figure 2A**). However, we did observe a sharp increase in reads mapping to chr14 within the cfDNA sample taken at the end of induction therapy (red dashed line, **Figure 2A**). This spike might be attributed to a clonal expansion of a B-cell caused by an illness or other antibody response in the patient. Just as with all other types of DNA sequencing used for ALL clonality detection, the major caveat of our MinION workflow is that it cannot distinguish between B-ALL and normal B-cells.

Like AAL-010, patient AAL-012 had a clone, IGHV4-39, not present in the diagnosis gDNA bone marrow biopsy sample but identified in the diagnosis cfDNA plasma sample (**Figure 2B**). Because every B-ALL clone releases cfDNA into the plasma, cfDNA may ultimately be more accurate in detecting the actual clonal composition of the B-ALL than the bone marrow biopsy, which can only detect clones present at the site of biopsy. Additionally, early into both patient AAL-10’s and AAL-012’s treatment, we observed a sharp increase in cfDNA associated with these clones, which likely indicates leukemia cell death and release of DNA into the blood and may indicate a good response to treatment.

Two additional patients, AAL-004 and AAL-008, had blasts present in their clinical samples at the end of induction therapy. Patient AAL-004 was not diagnosed with MRD, as the blast count fell below the one blast per 10,000 cell threshold for clinical MRD. Patient AAL-008 was MRD positive. In both patients, we detected heterogenous B-ALL comprised of several distinct IGHV sequences in both the gDNA from bone marrow biopsy and the plasma cfDNA isolated at diagnosis. Patient AAL-004 appeared to clear their clones during induction therapy, as we could not detect reads linked with chr14 in the cfDNA samples. However, we observed clone IGHV3-74, a major clone present at diagnosis, re-emerge in cfDNA samples collected at the end of consolidation therapy (**Figure 2C**). Patient AAL-008 unfortunately never cleared the clones detected in the biopsy gDNA and cfDNA sample at diagnosis. Interestingly, we observed that clone IGHV3-30, a minor clone in the cfDNA at diagnosis, was the predominant clone at the end of consolidation therapy (**Figure 2D**). Together, these data highlight the utility of cfDNA analysis as a method to track B-ALL clonal dynamics during patient treatment. This workflow may be useful in identifying patients whose B-ALL does not respond well to therapy, such as patient AAL-008, which may allow them to be stratified into a high-risk category at an early time point so that they can receive additional treatment. Additionally, cfDNA can be used as a non-invasive method to routinely assess patients for clone re-emergence after treatment.

### cfDNA can be detected in the CSF of B-ALL patients using the MinION sequencing workflow

Approximately 4% of B-ALL patients will have infiltration of ALL blasts into their CNS at the diagnosis, and 30-40% of the patients will have pronounced CNS disease at relapse (4). CNS infiltration is a significant adverse prognostic indicator in B-ALL, but the only way to diagnose it is to detect blasts in the CSF sample. However, B-ALL cells can be present in the brain or associated with spinal neurons and not be freely floating and detectable in the CSF, meaning patients might have CNS disease but present clinically as negative for CNS infiltration. For this reason, every ALL patient is given intrathecal chemotherapy. We reasoned that since cfDNA is released by B-ALL cells into patient biofluids, it might be detectable in CSF samples even if the blasts themselves are not. We applied our MinION sequencing workflow to cfDNA isolated from CSF samples taken from patients diagnosed as CNS positive (AAL-008 and 010) and CNS negative (AAL-009 and 012). As expected, we detected IGHV reads in the cfDNA from CNS positive B-ALL patients (**Figure 3A-B**), and these clones were cleared by the end of induction therapy. Interestingly, we detected very low levels of IGHV sequence in the diagnosis CSF cfDNA sample from patient AAL-009 (**Figure 3C**), even though this patient was diagnosed as CNS negative. Finally, patient AAL-012 was CNS negative at diagnosis. However, we observed an IGHV read in the cfDNA from the CSF of this patient at diagnosis that significantly expanded by the end of induction (**Figure 3D**). CSF is not drawn from patients after induction therapy if they are diagnosed as CNS negative, so we could not follow up on the IGHV in the cfDNA in later CSF samples of AAL-012. However, their plasma cfDNA remained clear of IGHV reads throughout consolidation. Finally, across all patients, we observed that some of the IGHV reads in the cfDNA from the plasma were also present in the cfDNA in the CSF, indicating B-ALL clones were present at both locations. In other instances, clones were exclusively present in one location, suggesting that cfDNA analysis may be useful in research focused on ALL homing to the CNS.

**Figure 3.**
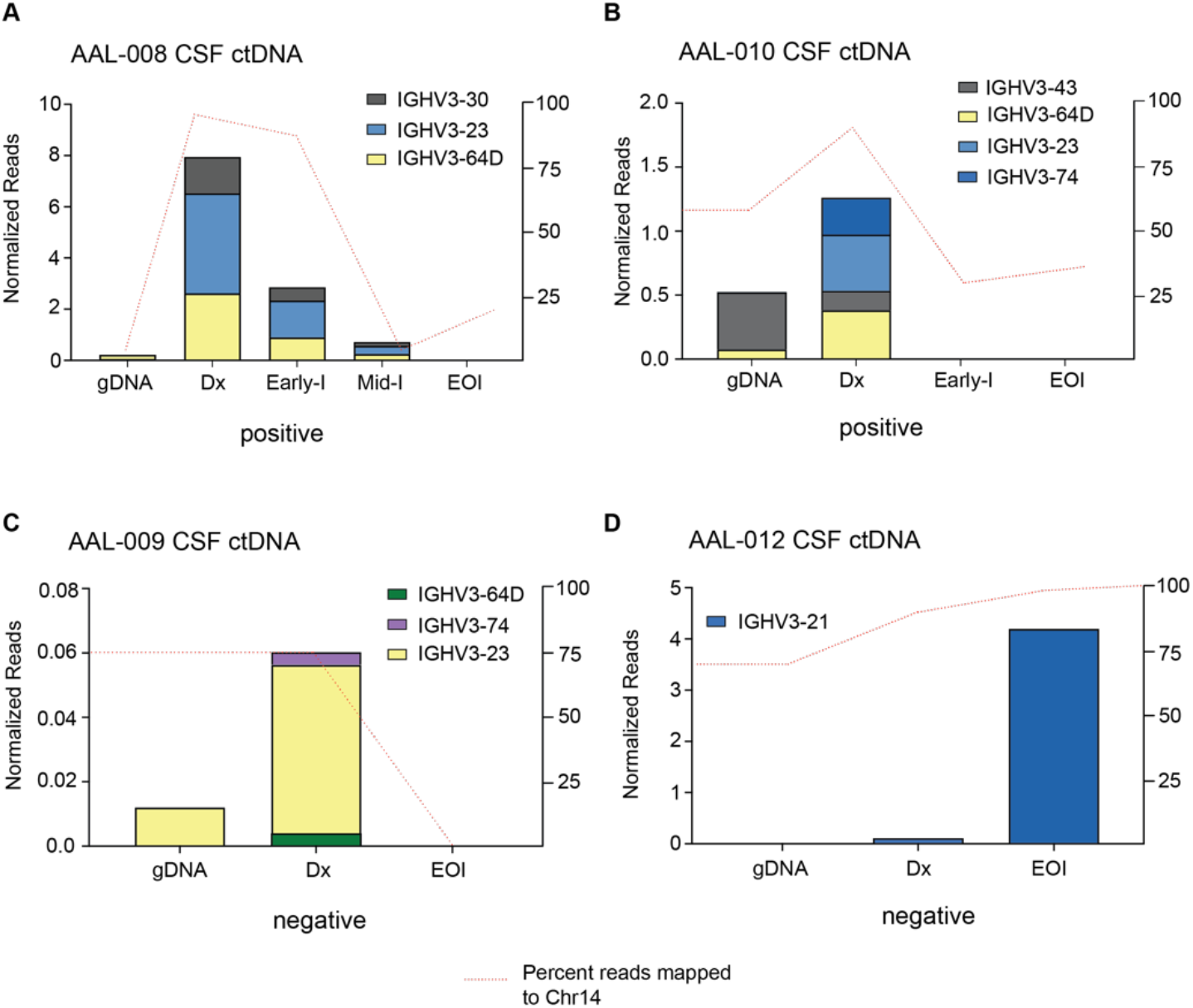
IGHV sequencing can detect cfDNA in the cerebrospinal fluid of B-ALL patients. cfDNA was extracted from CSF samples of four patients, IGHV regions were PCR amplified, and resultant libraries were sequenced using MinION. Data were analyzed using Galaxy. Clones in the genomic DNA diagnosis cell-free DNA (Dx) CSF samples that comprised at least 5% of total reads were tracked across all timepoints, and in genomic DNA isolated from bone marrow biopsy. Clone abundance is normalized to total mapped reads per library. Percent reads aligned to Chr14 (red line) is a relative indicator of leukocyte abundance at each timepoint/sample. Patients in (**A**) and (**B**) were clinically diagnosed as CNS positive, indicating infiltration of B-ALL cells into the CNS. Patient (**C**) and (**D**) were diagnosed as CNS negative, although IGHV reads were detectable in the CNS sample.

Our study demonstrates the utility of a nanopore sequencing workflow as a simple, rapid, and low-cost method to assess clonal VDJ rearrangements in B-ALL. This workflow will be useful to research laboratories interested in lymphocytic clonality; similar nanopore sequencing strategies could be developed and applied to detect tumor-associated mutations in cfDNA in any cancer type. We found that quantitation of IGHV reads from cfDNA samples provided insights into B-ALL heterogeneity and could be used to assess the dynamics of B-ALL clones in response to treatment. Our data suggest that cfDNA monitoring could be a useful non-invasive method to routinely monitor ALL burden in patients undergoing treatment, which may allow for earlier diagnosis of MRD and more accurate risk stratification. Analysis of cfDNA may also ultimately prove to be a clinically useful complement to the current cell-based ClonoSEQ methodology in diagnosing CNS disease, in which blasts may not be physically present in the CSF sample but are continuously releasing cfDNA.

## Methods

Further information can be found in the Supplemental Methods.

*Study approval*. All patient samples were collected after obtaining informed consent according to protocol 44672, approved by the University of Kentucky’s Institutional Review Board.

## Author Contributions

SS, YC, MGH, TCB, and JSB designed experiments; SS and YC conducted experiments and acquired data; SS, YC, IAS analyzed data; TJT, BLF, TCB consented patients and collected samples; MGH, LHM, AHC, IAS processed samples; SS, YC, MGH, and IAS drafted the manuscript; JSB edited the manuscript; JSB provided supervision and funding.

## Acknowledgments

Funding for this research was provided by a Kentucky Pediatric Cancer Research Trust Fund research grant to JSB and TCB, NIH grants DP2CA228043 and R37CA227656 to JSB, as well as T32CA165990 to MGH. The graphical abstract was created with Biorender.com.

