## Supplemental Figures and Methods for "Nanopore sequencing methods detect cell-free DNA associated with MRD and CNS infiltration in pediatric Acute Lymphoblastic Leukemia"

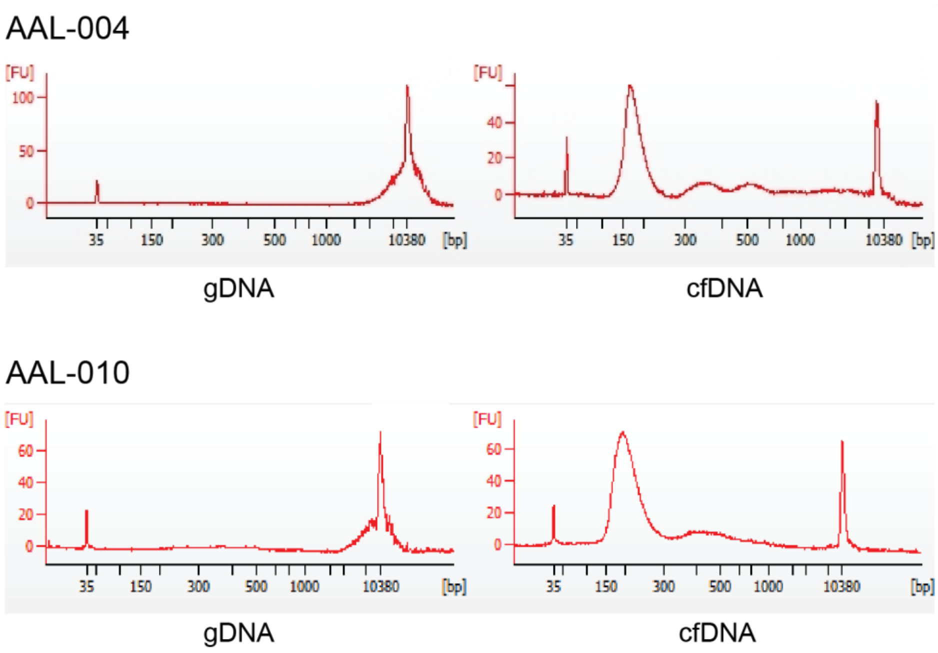


**Supplemental Figure 1. Electropherograms distinguish genomic DNA from cfDNA.** Genomic DNA (gDNA) and cell-free DNA(cfDNA) samples purified from the diagnosis bone marrow biopsy and blood draw from patients AAL-004 and AAL-010 were run in an Agilent Bioanalyzer 2100 using a high-sensitivity DNA kit. Genomic DNA appears as sizes >10,000bp (denoted on the X axis), while cfDNA appears as ~150bp peak. The y-axis denotes signal intensity.

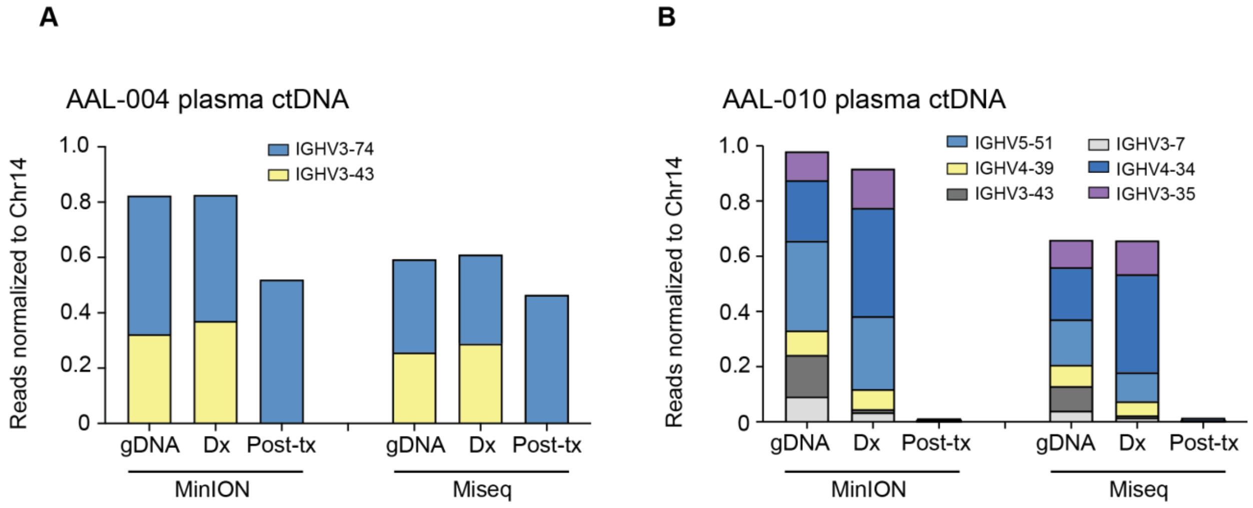


**Supplemental Figure 2. Comparison of clone distribution between Illumina and Nanopore MinION sequencing platforms**

IGHV PCR of select timepoints from patients AAL-004 (**A**) and AAL-010 (**B**) was divided and sequenced with MinION (nanopore) or MiSeq (Illumina). V-region clones comprising more than 5% of total clones in sample taken at diagnosis were tracked across the two later time points and normalized to the total number of reads that aligned to Chr14 per sample. Clone distribution across different timepoints within each patient is uniform regardless of sequencing platform. Dx=sample taken at B-ALL diagnosis, post-tx=sample taken post-treatment.

**Supplemental Methods**

*Isolation and cryopreservation of mononuclear cells from bone marrow aspirate.*

All patient samples were collected after obtaining informed consent according to protocol 44672, approved by the University of Kentucky's Institutional Review Board. Bone marrow aspirate was collected in Streck tubes (Streck, Cat # 218962) at the University of Kentucky's Pediatric Oncology clinic. Samples were diluted with an equal volume of room temperature 2% fetal bovine serum (FBS) in 1X PBS. The sample was slowly added to a 50mL Sepmate tube (Stem Cell Technologies, 85450), containing 15mL of Ficoll-Paque density gradient medium (Millipore Sigma, GE17-5446-52) and centrifuged at 1,400rpm for 10 min. The top layer containing the mononuclear cells was poured into a new 50mL conical tube. The volume was brought up to 50mL with RPMI supplemented with 10% FBS and spun for 15 mins at 1,400 rpm at 4°C. Media was aspirated, cells were resuspended in 1mL RPMI, then counted with a hemocytometer. An additional 49 mL of complete RPMI was added to the cells, and cells were centrifuged for additional 10 mins at 1,400 rpm at 4°C for the second wash. The media was removed, and the cells were resuspended in 1ml of freezing medium (90% FBS + 10% DMSO) per $10^7$ cells and added to a cryovial. Vials were placed in a CoolCell (Corning, 432003), cooled at -1°C/minute at -80°C then moved to liquid nitrogen for long-term storage.

*Isolation of genomic DNA from banked mononuclear cells.*

Vials of banked mononuclear cells from patient bone marrow aspirate were warmed to 37°C and transferred to a sterile 15 mL conical tube. Pre-warmed, 25% fetal bovine serum (FBS) in Iscove's Modified Dulbecco's Media (IMDM) was added dropwise to the cells at a rate of approximately 2-3 seconds/ml for 10 mL. Cells were centrifuged at 1,000 rpm for 10 minutes, and the supernatant was gently aspirated. The addition of media, centrifugation, and removal of supernatant was repeated. Cells were then washed once in 1X PBS. gDNA was isolated using the Zymo Quick-gDNA Miniprep Kit (Zymo, D3025), using the manufacturer's instructions.

*Isolation of cfDNA from Plasma and Cerebrospinal Fluid (CSF)*

Patient blood and CSF samples were collected into cell-free DNA collection tubes (Streck, 218962). Samples were stored at room temperature and processed within two weeks of collection. Blood samples were centrifuged at room temperature for 10 minutes at 1,600xg. After the initial spin, the plasma was removed, being careful not to disturb the buffy coat layer, and transferred into a clean 1.5 mL Eppendorf tube, and centrifuged for 10 mins at 16,000xg at room temperature. The supernatant containing the cfDNA was carefully removed so as not to disturb the cell pellet and placed in a fresh tube. cfDNA from the plasma was isolated using the Qiagen QIAmp MinElute ccfDNA Midi Kit (Qiagen, 55284) as per the manufacturer's instructions, except that the final elution occurs with nuclease-free water pre-warmed at 56°C for an enhanced yield of DNA from the column. cfDNA was isolated from CSF samples using the Zymo Research Quick-cfDNA Serum and Plasma Kit (Zymo, D4076) according to the manufacturer's instructions except with the following modifications: i) after adding proteinase K, the samples were incubated at 55°C for 1hr and ii) final elution step was carried out in 35 µl of nuclease-free water pre-warmed at 56°C. cfDNA samples were quantified using a Qubit Fluorometer using the high-sensitivity dsDNA quantification kit (Thermofisher Scientific, Q32851) according to the manufacturer's instructions and stored at -20°C.

*PCR amplification of IGH regions in genomic and cell-free DNA*

The IdentiClone *IGH* Gene Clonality gel detection kit (Invivoscribe, 91010020) was used to amplify the *IGH* region of genomic and cfDNA. The PCR was performed using the primer master mix labeled Tube A (Invivoscribe, 21010010CE) according to the manufacturer's directions, with 0.5ng of genomic DNA or cfDNA as input. Amplitaq Gold Taq Polymerase (ThermoFisher Scientific, N8080240) was used for PCR amplification at 0.2ul per 50ul reaction, and PCR was run for 40 cycles with the cycling parameters of 95℃ for 7 minutes of initial denaturation; 40 cycles of 95℃ for 45 sec; 60℃ for 45 sec; 72℃ for 90 sec and final extension of 72℃ for 10min. A portion (5µl) of the PCR reaction was run on agarose gel electrophoresis to confirm the amplification before proceeding with the library preparation.

*Library preparation for MinION sequencing*

Following the *IGH* clonality, PCR samples were measured on Qubit, and 50 ng of each sample was used as input for the library prep for the MinION sequencing. The PCR Barcoding Kit (Oxford, SQK-PBK004) was used for the library generation according to the manufacturer's protocol with the following modifications: i) since an amplicon was used as starting material, the initial fragmentation step of the protocol was omitted, ii) half-reaction volumes were used throughout the library generation procedure to conserve materials. Libraries were loaded onto the MinION Nanopore sequencer (Oxford) and were run until all barcodes had a minimum of 4000 reads, which averaged about 1-2 hours.

*Illumina MiSeq analysis*

*IGH* variable region was amplified from plasma cfDNA using the Invivoscribe kit as described above. The PCR reaction was size-selected using Ampure bead purification and eluted in nuclease-free water. The eluted DNA was sent to Genewiz for targeted amplicon-based sequencing under their "Amplicon EZ" category. Standard Illumina adapters were used in the library prep, and the final amplicon libraries were run on the Illumina MiSeq sequencing system. The merged Fastq files were then analyzed similarly to the files from the MinION, described below, to assess the IGH variable region clonal rearrangements and their abundance.

*MinION analysis pipeline to identify IgH rearrangements*

Sequencing reads were base-called and demultiplexed using the MinKNOW software package and built-in basecaller. Reads with a quality score of ≥ 7 were used to generate fastq files and for all downstream analyses. Fastq output files were concatenated into a single file per sample. Merged fastq files from MinION and those generated by MiSeq were then processed using the Galaxy server and its available tools as follows. Reads were mapped to hg38 using Minimap2 to generate BAM alignment files [1]. Alignment files were then used as input for Feature Counts along with a .gff reference file of the IGHV region on chromosome 14. The output text file of how many reads mapped to each feature in the .gff reference file was used for data analysis and graph creation. Read counts for each feature were normalized as a percentage of total reads for each patient. Major IGHV clones were identified in the diagnosis sample as those features that contained reads equal to or greater than 5% of the total reads for that sample. The select major clones were tracked through subsequent time points in each patient. All graphs were generated using Prism GraphPad.

***Extended Methods for Nanopore Sequence Analysis***

**Software Packages Required:**

1. Account with Galaxy (usegalaxy.org)

2. Minimap2 (hosted by Galaxy)

3. FeatureCounts (hosted by Galaxy)

**Input Files Required**:

1. .fastq read files from a sequencing experiment. These should be one file per experiment. If experiment generates multiple files, merge them into one before proceeding.

2. hg38.IGHV.gff3 feature file. This file annotates the different genomic features in the IGHV region (i.e., genes, exons, etc) and their corresponding genomic positions (i.e., Chr14:101,224-115-032). It was created by trimming the Chr14.gff3 file down to the IGHV region.

**Analysis Protocol**:

**<u>A. Minimap2</u>**

This software package is used for mapping sequencing data to a reference genome to determine read locations and sequencing accuracy. This particular mapper is optimized for handling long reads, but can just as easily align short-read data.

1. **Log into Galaxy and use the "Upload Data" icon to upload the two input files onto the Galaxy server.** A new window will pop up where you can select the files of interest from a local computer or drag-and-drop them in. Press "start" to begin upload. Correctly uploaded files will appear as green tabs on the left-hand side of the screen.

2. **Search for Minimap2 software by typing it into the search bar in the top right corner. Select the "map with minimpa2" option from the results**. The main window will now show a

graphical user interface for the minimap2 software package which will already have most fields populated with default presets. (Only change presents outlined below.)

3. **Select a Reference Genome**. In the drop-down menu of "Using a reference genome" select "Human (Homo sapiens) (b38):hg38". This is the most recent version of the human reference genome that you will use to map your reads.

4. **Select input file for mapping**. In the "Select fastq dataset" drop-down menu tab pick your .fastq input file. (If the file does not show up in the drop-down menu click the open file folder icon to the right of it to select it manually).

5. **Select Mapping Parameters**. In the "Select a profile preset options" drop-down menu pick the third option of Oxford Nanopore read to reference mapping.

6. **Select output format**. Expand the "Set advanced output options" tab and double check that the output format is set to BAM (not CRAM or PAF).

7. **Press Execute to run job**. The job will appear in the right-hand column as grey while it's in the queue and will turn pink when it begins processing. Once it is done it should turn green. This BAM output job will have two files associated with it: a .bam alignment file and a .bai index file. It will also serve as the input for the next step.

**B: FeatureCounts**:

This software tool counts the number of reads that are aligned at a specific annotated feature. It requires a read alignment file (BAM) and a feature reference file (gff or gff3) that it cross references against one another.

1. **Search for FeatureCounts software** by typing it into the search bar in the top right corner and clicking on the name in the results.

2**. Select your alignment file.** In the "Alignment file" drop-down menu pick the Minimap2 alignment file from above. (If the file does not show up in the drop-down menu click the open file folder icon to the right of it to select it manually).

3. **Select the Gene Annotation file**. In the "Gene Annotation File" drop-down menu pick the "in your history" option and then select the hg38.IGHV.gff3 file you uploaded earlier in the space that appears.

4. **Specify counting parameters**. Expand the "Advanced options" tab. Here you will specify which features you want to count since a gff file will contain many feature types. In the "GFF feature type filter" put "gene" and in "GFF gene identifier" put "Name". Case is IMPORTANT here since software searches for exact matches. Scroll down to "Long Reads" option and turn it to Yes.

5. **Press Execute to run job**. Job will appear in the right-hand column. The output will generate two files. One will be the actual counts at every feature selected from the gff file and the second will be a Summary file of read statistics.
